# Rapid active-reflection display on dorsal stripes of *Atherinomorus lacunosus*

**DOI:** 10.1101/2021.02.15.431186

**Authors:** Masakazu Iwasaka

**Author notes:** Correspondence to: Masakazu Iwasaka.

## Abstract

A large number of living creatures are able to use environmental light effectively as a biological display. The biological structural colors are very attractive not only within the coloring species but also to humans. However, the detailed function of bio-reflectors, which constitute the structural color with respect to communication, remains unknown. *Atherinomorus lacunosus* has alignments of iridophore spots on its dorsal part. Here it is found that a spot with a diameter of approximately 0.1 mm causes a rhythmic blinking of light owing to rapid reflection changes in iridophores existing inside the spot. The iridophores contain reflecting particles which show similar rotational responses to magnetic field under a light exposure. The speed of the intensity change of light at a frequency of approximately 1 Hz is proposed to be controlled by the nervous system of *A. lacunosus*. This kind of passive illumination may contribute to the development of a new optical device with low energy consumption.

## 1. Introduction

Studies on cephalopods, cuttlefish [1–3] and octopus [4] have revealed that a dynamic camouflage effect occurs in the cellular tissue existing near the skin. A similar mechanism for coloring changes has also been discovered in the tissue structures of fish, [5–10] insects [11–16] and a mammal [17]. In biological materials that are able to control light, guanine is useful for light reflection and optical interference in iridescent tissues. [5–7] Guanine-based photonic crystal structures have been found in many species of fish, [5,6,10,18–21] scallop [22] and animal plankton. [23,24] It is often possible to observe many markings on the surface of a fish body, although their purpose in the life of the fish remains to be clarified. For example, the side part and dorsal part of a fish are frequently colored in silver and black, respectively. The color is generated by several kinds of chromatophores. [25]

It has also been indicated that the color patterning by several kinds of chromatophores containing a pigment is concerned with both hormonal and neural activity. [26] Additionally, the role of patterns on a fish body surface has been suggested to be an active communication tool, while nonlinear mechanism studies have imputed the stripe pattern in a fish to a Turing pattern, which was generated by an interaction between the chromatophores. [27,28,29,30]

Developmental pathways, possibly controlled by the interactions between the same and different kinds of chromatophores, are considered to be an interesting and important topic. The effects of cell-to-cell interactions on the color pattern formation in zebrafish have been investigated in recent studies. [31–33]

The silvery shine of the side and ventral parts is mainly controlled by iridophores containing guanine crystals. Over the dorsal region that is mainly covered by black melanophores, some species, such as Japanese anchovy, have an additional layer that looks partially blue to purple. We conjecture that the additional color modification is carried out by light interference in guanine crystal layers. Detailed analyses of the fish coloring pattern have been performed on the skin of zebrafish, in which a variety of patterns, stripes and spots have been inspected from the viewpoints of developmental and ethological features. [34]

Most cases of coloring on a fish body are static, that is, non-dynamic coloring. Black coloring of the dorsal part of a fish is explained by a counter shading effect, which helps the fish become less visible to a predator approaching from above. In contrast, the ventral part covered with shiny iridophores can result in a matching of the light intensity between the fish body and water for a predator looking from below. As the guanine crystals that result in light reflection are fixed behind the scales, the skin tissue itself performs a passive reflection. Control of the light reflection can be carried out by the swimming motion of the fish.

A species of fish has a light emitting tissue—a photophore—in the ventral or side part of the body, and the photophores that are distributed over the surface of the ventral to side parts exhibit an active matching of the light intensity, known as counter illumination. The role of reflector particles, such as guanine crystals, still requires further clarification.

The present study focuses on how the reflecting spots behave like a light switching panel on the dorsal skin of *Atherinomorus lacunosus*. *A. lacunosus* is one of the genera belonging to silversides, order *Atheriniformes*. For the first time, an active and rhythmic light reflection on fish skin is discovered. The reflecting particles existing inside the spot exhibit a rapid change of brilliance at approximately 1 Hz for one-directional light irradiation.

Previous studies have reported that some species of squid show a dynamic structural color change. It has been suggested that the color change is controlled by the neural system of the squid.

However, the time to obtain a sufficient color change is more than several seconds. Studies on neon tetra have also provided data about its dynamic color control in chromatophores on a similar time scale. In the case of a chameleon, the color change speed is of the same order. Here, we present the quickest change of color or light intensity in a species of silversides.

## 2. Materials and Methods

### 2.1 Sample and image analysis

Specimens of *Atherinomorus lacunosus* were obtained in Okinawa. Animal experiments in this study were carried out in accordance with the policies of Hiroshima University Animal Care and Use Committee (approval number: F19-2, Hiroshima University).

### 2.2 Image analysis

The iridophore spots on the body surface were analyzed by using digital image processing units. Low resolution images of the fish body surface were taken by using a digital camera (Coolpix AW130, NIKON co. Tokyo, Japan) and a digital microscope (PC-230, Vixen, Tokyo, Japan). For the high-resolution images with an amplification of ×230, the same digital microscope was used. The images of fish and its body surface were obtained under exposure to white LED light (MLEP-B070W1LRD, MCEP-CW8-2M, Moritex, Tokyo, Japan) whose light polarization was not additionally modulated. The light illumination was provided from the side and the scattered or reflected light from the sample was monitored. The incident angle of the irradiation was 50–80 degrees. Additionally, by changing the direction of the irradiation on the plane of the observed area, images with the highest and lowest contrast were captured. The observation of the sample was carried out under a wet condition, where the surface of the sample was soaked in water containing anesthetic solution (2-phenoxy-ethanol).

Image analyses were performed using Image J (NIH) software. Radiance of the sampled image was measured and quantified by the gray scale value, which was produced from the intensities of red (R), green (G) and blue (B) pixels. When green color of an image was suitable for evaluation, the gray scale value of B was employed.

### 2.3 Mimicry of iridophore spots

A realistic model of the iridophore spot was formed on a transparent acrylic plate whose size was 20 mm × 20 mm in the plane and 1 mm in thickness. An aqueous suspension containing guanine platelets was facilitated in a cylindrical well, which was fabricated in the plate. The well, approximately 1.2 mm in diameter, was bored by an end mill of a router. Guanine platelets of sea bream, *Sparidae* were suspended in water and were poured into the well, and the micro-platelets were floating in the water during the observation with and without magnetic field exposure.

The fabricated acrylic plate was placed on the gap space between magnetic poles of a resistive electromagnet (WS15-40-5K-MS, Hayama Inc., Koriyama, Japan). The gap length in the magnetic field exposure space with a maximum field of 500 mT was 70 mm. Applying the magnetic fields, brilliance changes in the formed spot were recorded by a digital microscope system (RH-2000 with MXB-2500REZ (lens), Hirox co., Tokyo, Japan) under one-directional light irradiation from the side of the plate. LED light (LA-HDF158A, Hayashi-Repic, co. ltd, Tokyo, Japan) was used as the light source.

## 3. Results and Discussion

A photograph of the belt-like pattern in the dorsal part of *A. lacunosus* is shown in Figure 1. This belt consisted of a wave-like alignment of chromatophores. Magnifications of the image (Fig. 1 *(b)*) show circular spots that were aligned along the edge of a scale. The line of circular spots, which was composed of connecting spots, extended from the ventral to dorsal sides. This iridophore-like spot overlapped with a black melanophore which had a dendrite. In Fig. 1*(a)* and *(b)*, silver and green long bands were observed on the side of the body of the fish. In addition to the structural color, green, blue and yellow colors were observed in the dorsal belt and also in the parietal skin.

**Figure 1.**
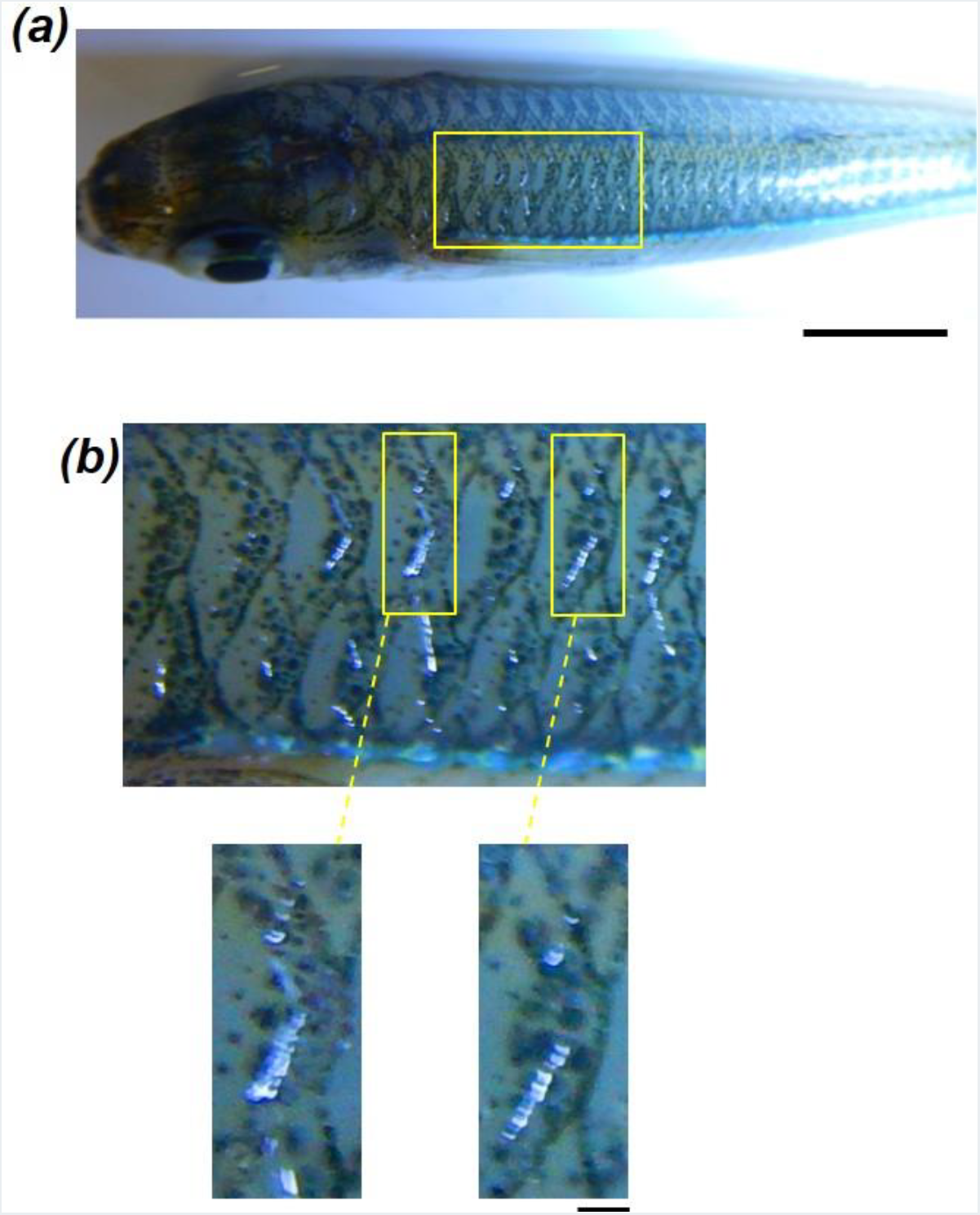
Reflecting spot arrays on the dorsal part of *Atherinomorus lacunosus*. *(a)* Top view of the dorsal part showing belts of spots aligned along the edges of a scale. Bar is 10 mm. *(b)* Magnified images of the belts. Bar is 0.5 mm.

Exposing this iridophore-like spot to light irradiation along the optical axis of the lens showed no remarkable reflection change. However, irradiating light from a side direction exhibited a dynamic brilliance change, as shown in Figure 2. In the image shown in Fig. 2*(a)*, the fringe of the circular spot suddenly became brilliant when side illumination was supplied from the right side. Inside the spot, the green and yellow particles became distinct.

**Figure 2.**
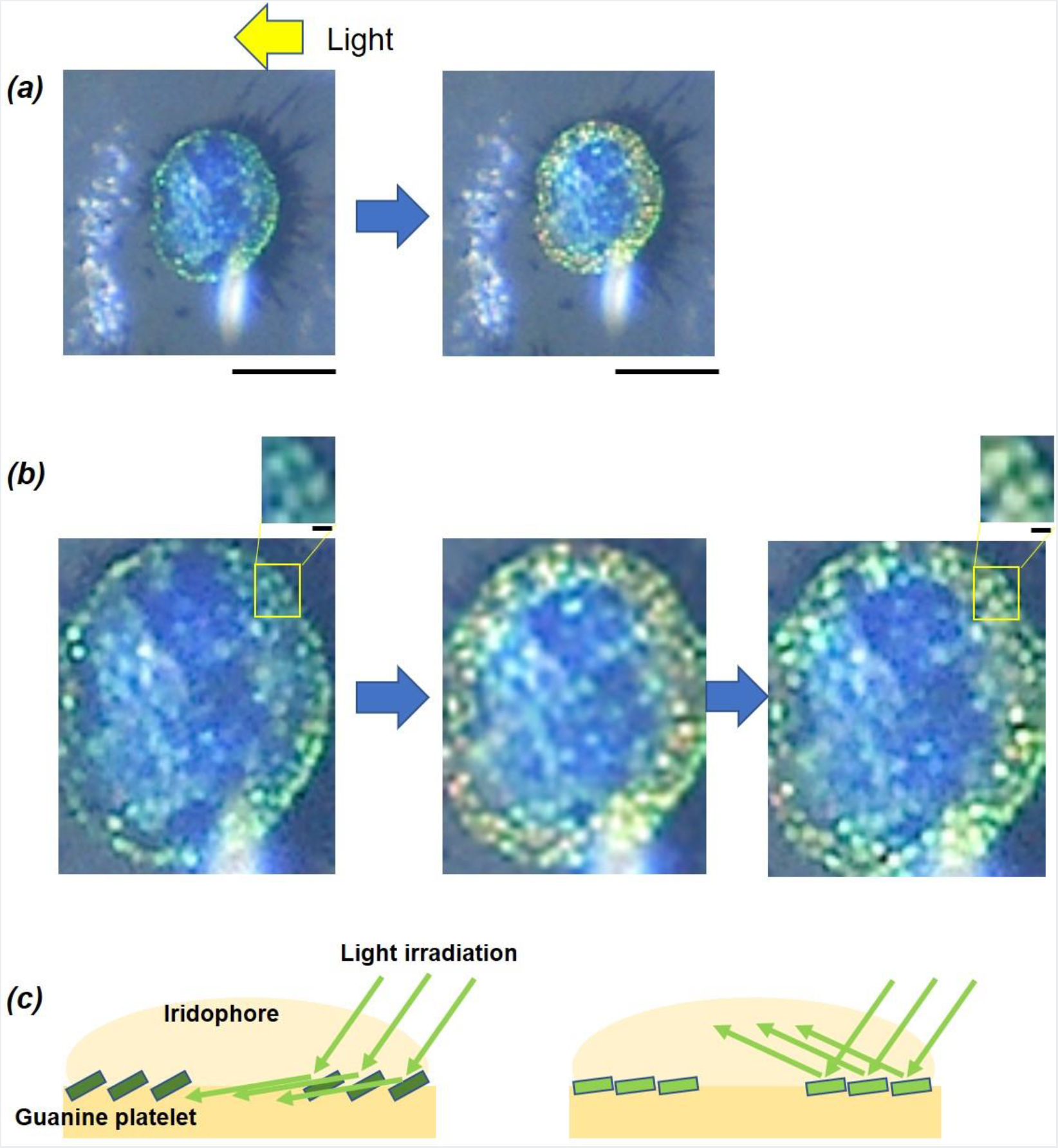
Reflection enhancement in an iridescent spot under side light illumination. *(a)* Change of brilliance in the fringe of the circular spot. Scale bar is 100 mm. *(b)* Expanded images of the spot containing micro particles. Upper small cuts show four particles that resemble platelets, probably causing an inclined surface. Scale bar is 5 μm. *(c)* Illustrations of the iridophore. The reflector platelet, assumed to be made of guanine, is controlling the direction of light reflection.

A magnification of the spot is shown in Figure 2*(b)*, where it was obvious that active light reflection of this fish was achieved by micro particles. The size of the circular reflecting particles was 5–10 micrometers in diameter, which is a similar size as the guanine crystal platelets from a fish, for example, sea bream. Behind the iridophore, a pseudopod was observed within the penetration of the used lens. This suggested that the circular shape of the reflecting particle was the broad surface of a platelet rather than the cross-section of a fiber elongated to the vision.

The brilliance became dark when the light irradiation was terminated. This proved that the brilliance of the spot arose from light reflection and not light emission such as bioluminescence. Additionally, the changes in brilliance and colors when the angle of incident light was varied exactly resembled those of accumulated guanine platelets from other fishes. A platelet or a stack of platelets were aligned in a spot. When a trigger for changing the incline of the platelets by any mechanical force occurred, the reflector platelets of the peripheral part oriented themselves to enhance the brilliance.

A tunable color change in biological tissue has been reported previously in cuttlefish and neon tetra. These studies focused on the change in thickness and angle of the layers, which control the optical interference that dominates its structural color. The newly discovered feature in spots of *A. lacunosus* suggests the possibility of a rapid response in reflection only. This study found for the first time that the spots caused a rapid blinking of the reflected light. Figure 3*(a)* shows an example of the reflection blinking in connected spots formed a line. The analysis on a time course of the green light reflection intensity revealed the occurrence of synchronized light blinking by three spots in the line. This blinking occasionally repeated with a rhythm like respiration. Another type of arrangement of spots, scattered spots, is shown in Figure 3*(b)*. There was no distinct synchronization of individual blinking. In the scattered spots area, the light reflection changes of the spots occurred separately.

**Figure 3.**
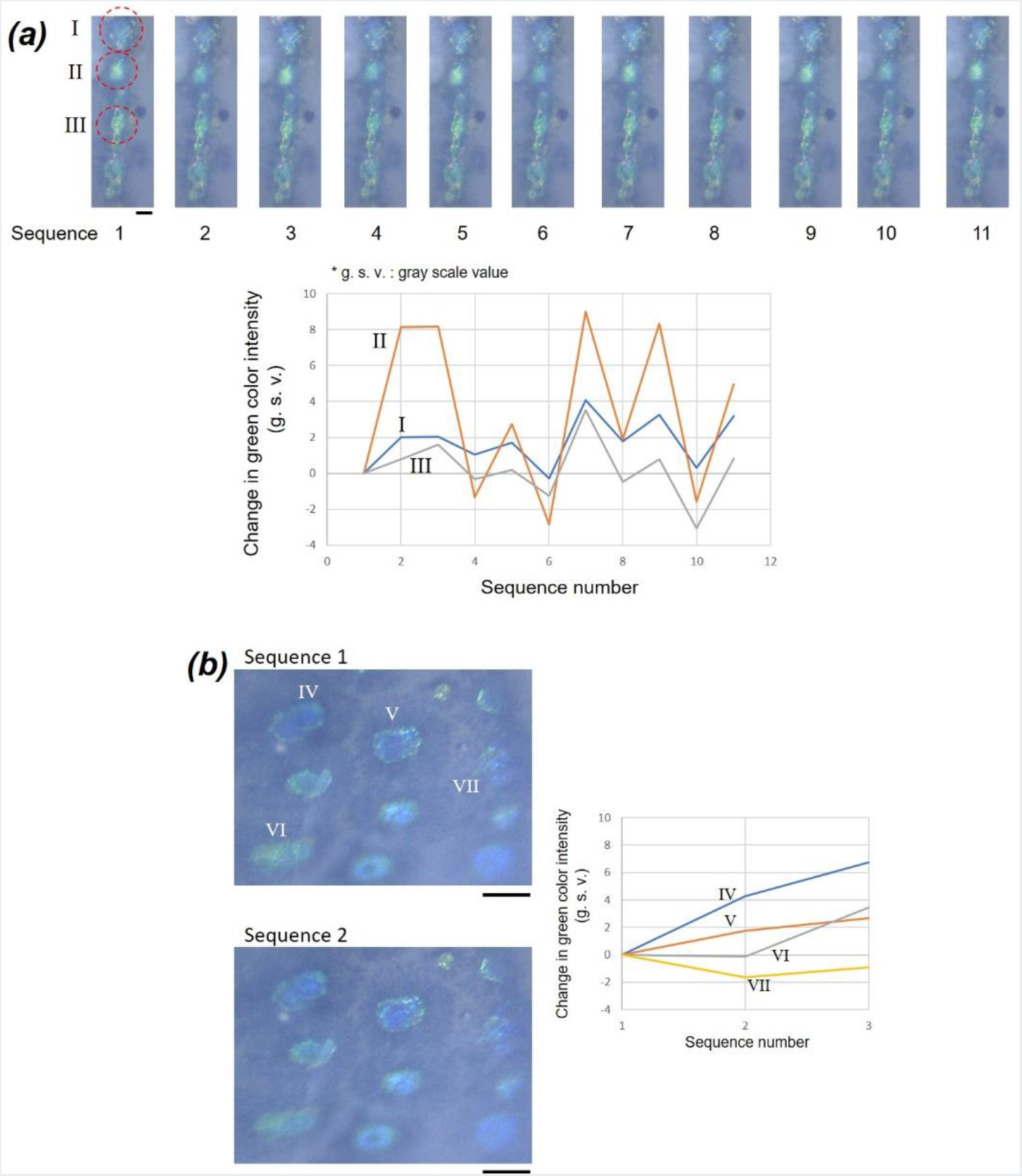
Blinking iridescent spots with synchronization *(a)* and desynchronization *(b)*. *(a)* Synchronized blinking in three spots (I–III) in a group of connected spots forming a line. Length of time between two sequences is approximately 0.5–1.0 s. Brilliance (gray scale value, g.s.v.) of green color only was calculated. *(b)* Desynchronized blinking in three spots (IV–VII) in a group of scattered spots. Scale bars are 100 μm.

The origin of blinking is ascertained in the following discussion on a careful test of the correlation between influencing factors. First, less of an effect of the skin motion on the reflection changes in the spot exists. Although a discontinuity of the refractive index of the scale and water can change the incident angle of light when the scales slide owing to skin motion, no correlation was found between the spot reflection and the sliding.

It was apparent that the reflector particles in the spot had light reflection anisotropy resembling that of guanine crystal platelets existing in fish skin and choroidal tapetum. The typical size of a fish guanine platelet was a few micrometers to several tens of micrometers in length and around 100 nm in thickness. [18–21, 35] Therefore, it is possible to mimic the reflecting spot of *A. lacunosus* by using guanine platelets extracted from a skin of fish, such as goldfish.

Figure 4 shows a simulation of the spot formed in a hole in an acrylic plate. The aqueous solution containing guanine platelets of sea bream was localized in the hole. A demonstration of the light reflection change in the mimicked spot was carried out by applying a magnetic field at 500 mT. The mechanism of the magnetic orientation of guanine platelets owing to its diamagnetic anisotropy provided the inclination of the guanine-based reflector platelets floating in water. As a result of restraining the rotational axis of a platelet by applying the magnetic fields orthogonal to the incident light, an enhancement in the platelet that was similar to the brilliance changes shown in Fig. 2 was achieved.

**Figure 4.**
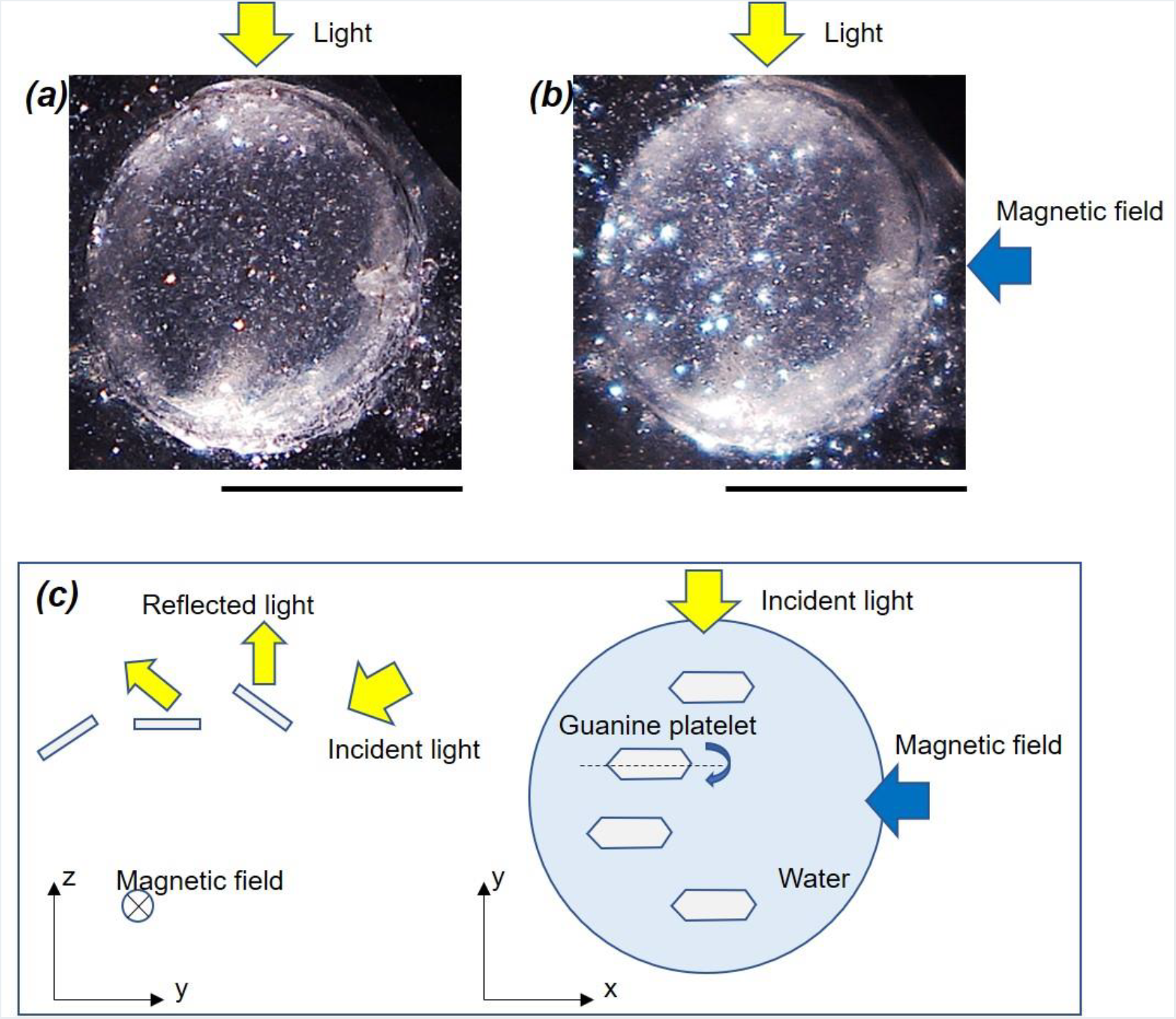
Experimental simulation of the blinking spot by using an aqueous solution containing guanine platelets of sea bream, *Sparidae* localizing in a hole formed in an acrylic plate. *(a)* Floating guanine platelets in a well containing water. Side light illumination is provided from the bottom. *(b)* Under exposure of a magnetic field at 500 mT. A, B; Scale bar is 1 mm. *(c)* Mechanism of the magnetically-induced incline change in guanine platelet and the light reflection modulation.

One of the characteristic colors that is produced by periodically aligned guanine platelets is blue. The coloring can result from the relative distance and angle between platelets. The mechanism for controlling the brilliance in spots of the dorsal belt of *A. lacunosus* within one second should be determined by the intracellular mechanical parts controlling the spacing and tilting angle of the reflector platelets. It was proposed that actin protein is one of the candidates that controls active color tuning in an iridophore [26]. It is questionable if actin fibers are able to perform a motion within one second inside the cell. Past studies have suggested the motion of tunable coloring on longer time spans. As a mechanical transducer tilts the platelet incline, liquid pressure actuation may be involved. A recent study found the effect of osmotic pressure on blue coloring in a photophore of luminous lanternfish.

Compared with previous findings, the repeated blinking in a reflecting spot of *A. lacunosus* resulting in a faster light intensity change with a frequency of ~1 Hz suggested the direct government of an iridophore by the nervous system. The speed of the color change in squid should be influenced by both body motion and neural signals controlling their iridophore. [3] It seems that the speed of reflection intensity changes in *A. lacunosus* was faster. Additionally, different spatial resolutions and key materials of the reflector unit were used for each species. The reflector platelets of *A. lacunosus* are compactly packed in the circular spot with a diameter of 0.1 mm. This may suggest that the use of this spot contributes to a short distance communication between fishes whose body length is ~100 mm.

There is a possibility that the light reflection involves exited light from an unknown fluorescence molecule. Some literature has reported the optical properties of a guanine crystal of fish from the viewpoint of fluorescence. A complete explanation of the optics in a guanine platelet of fish is not possible at present; however, the rapid light intensity change in the spot reflection requires some mechanical apparatus that is driven by a neural signal. The aim of this apparatus should be the adjustment of light intensity and color balance for the efficient display of the back of *A. lacunosus*. The belt of spots resembles an illumination board for inter-species communication, which remains for bio-ethological research to clarify the meaning of forming a dynamic pattern of light spots on a two-dimensional display by reflecting environmental light. The analysis of the biological optical tool can provide us with an opportunity to make a mechanical actuator of micrometer-sized optical parts embedded in a wet biological tissue.

Circular spot alignments in dorsal and parietal skin of *A. lacunosus* realize a blinking that resembles an apparent movement. The biological tissue constituted of reflecting micro-particles is actuated with a frequency of 1 Hz or higher.

## Authors’ contributions

M.I. performed the experimental design, experiments, measurements and analyses. All part of manuscript and illustrations were prepared by M. I.

## Acknowledgements

The author thanks Y. Uehara for collecting fish.

